# Sainsc: a computational tool for segmentation-free analysis of *in-situ* capture

**DOI:** 10.1101/2024.08.02.603879

**Authors:** Niklas Müller-Bötticher, Sebastian Tiesmeyer, Roland Eils, Naveed Ishaque

## Abstract

Spatially resolved transcriptomics has become the method of choice to characterise the complexity of biomedical tissue samples. Until recently, scientists have been restricted to profiling methods with high spatial resolution but for a limited set of genes or methods that can profile transcriptome-wide but at low spatial resolution. Through recent developments, there are now methods which offer subcellular spatial resolution and full transcriptome coverage. However, utilizing the high spatial and gene resolution of these new methods remains elusive due to several factors including low detection efficiency, high computational cost and difficulties in delineating cell borders. Here we present Sainsc (Segmentation-free analysis of *in-situ* capture data), which combines a cell-segmentation free approach with efficient data processing of transcriptome-wide nanometer resolution spatial data. Sainsc can generate cell-type maps with accurate cell-type assignment at a subcellular level, together with corresponding maps of the assignment scores that facilitate the interpretation in the local confidence of cell-type assignment. We demonstrate its utility and accuracy across different tissues and profiling methods. Compared to other methods, Sainsc requires lower computational resources and has scalable performance, enabling interactive data exploration. Sainsc is compatible with common data analysis frameworks and is available as open-source software in multiple programming languages.

## Introduction

Spatially resolved transcriptomics is becoming increasingly popular to characterise the complexity of biomedical tissue samples^1^. This cutting-edge technology allows researchers to map gene expression within the spatial context of tissue architecture, providing unprecedented insights into cellular heterogeneity and tissue organization^2^. Unlike single-cell RNA sequencing, spatial transcriptomics preserves spatial information, enabling the identification of cellular niches and the study of interactions between different cell types within their native microenvironment. This spatial context is crucial for understanding complex biological processes such as development, disease progression, and response to therapy^3^.

SRT methods can be broadly categorized into two groups^4,2^. The first are commonly referred to as imaging-based methods and offer single-molecule resolution through *in-situ* hybridization (ISH)^5,6^ or *in-situ* sequencing (ISS)^7,8^ for a selected number of target genes, normally in the range of 100s-1000s. The second are referred to as *in-situ* capture methods and incorporate spatial barcodes onto transcripts before sequencing, allowing whole transcriptome coverage but at limited spatial resolution^9,10^ (e.g. 100 μm inter-spot distance for Visium). The low spatial resolution of *in-situ* capture methods complicates the spatial analysis of single-cells, requiring deconvolution, imputation and/or integration with external single-cell transcriptomics resources^11–13^. However, recent advancements in spatial transcriptomics have revolutionized the field by offering full transcriptome profiling at nanometer resolution through profiling methods such as Stereo-seq, Seq-Scope, Open-ST, and Nova-ST^14–17^. These high-resolution techniques provide unique benefits, such as the ability to resolve transcriptome-wide expression at subcellular levels, in some cases in the sub-micron range.

However, these advancements also present several challenges. The sheer volume of data generated by these high-resolution methods requires robust and scalable computational tools for efficient data processing and analysis. Furthermore, measured gene expression is sparse, which could potentially arise from a combination of low capture efficiency and a smaller total capture area compared to lower resolution methods^18^. Furthermore, in the previous generation of supra-cellular spatial resolution *in-situ* capture SRTs, deconvolution of cell types was a common task, but with the increased sub-cellular spatial resolution of newer methods there is the potential to assign gene expression at the single cell level^19^. However, accurate cell segmentation remains elusive^20^, thus making correct assignment of measured transcripts to individual cells a non-trivial task. Addressing these computational challenges is essential for the widespread adoption and effective utilization of high-resolution spatial transcriptomics in biomedical research.

To address the current computational challenges in processing and analysing data from these methods, we developed Sainsc (pronounced “science”, segmentation-free analysis of *in-situ* capture data), which combines a cell-segmentation-free approach with efficient data processing of transcriptome-wide nanometer resolution SRT data. Sainsc is tightly integrated with common data structures and frameworks for single-cell and spatial data analysis in Python, rendering it highly accessible, while outsourcing the computationally demanding tasks to the Rust programming language enabling efficient and scalable computational performance in analysing full transcriptome nanometer scale data in native resolution. Furthermore, it’s segmentation free approach builds upon a Kernel Density Estimation (KDE) of gene expression, which reduces data sparsity and is suitable for classification tasks^21^.

We apply Sainsc to SRT data of several tissues, identifying cells missed by segmentation-based analysis, show practical applications of spatial maps of the assignment score to explain the confidence of the model results, and demonstrate generalisability across profiling methods. Furthermore, Sainsc shows significant computational performance advantages over other competing tools, therefore facilitating more efficient exploratory data analysis. Our implementation adheres to common data formats and analysis frameworks in the field to ensure interoperability, and the code is openly available on GitHub as Python and Julia packages.

## Results

### Sainsc – a computation tool for segmentation-free analysis of *in-situ* capture data

The Sainsc workflow is inspired by the SSAM and SSAM-lite algorithms^19,22^. Briefly, SSAM models spatial gene expression as a density through the application of Kernel Density Estimation (KDE), which can then be used for robust and accurate cell-type assignment across the tissue. The computational steps can be broadly broken down into reading data, quality control, data pre-processing, modelling spatial gene expression, generating or acquiring cell-type-specific gene expression patterns, and assignment of cell types to pixels to generate a cell-type map (**Figure 1**). Sainsc was developed to support efficient processing of *in-situ* capture SRT methods at their native resolution but can also be used to process data from imaging-based SRT methods through binning detected transcripts.

Sainsc primary implementation is in Python and leans on Rust for optimized data structures, data processing, and safe multi-threading support. The Rust elements enable computational efficiency, while the Python wrapper (using PyO3 for interfacing with Rust) enables accessibility to most computational life scientists. The main workflow is wrapped with a Python API and is provided with a broad range of convenience and plotting functions. Furthermore, we provide a Julia implementation of Sainsc’s main functionality given its rising popularity in the bioinformatics community^23^. To ensure interoperability with the wider ecosystem of spatial and single-cell analysis tools, Sainsc provides native support for reading established file formats (e.g. GEM files) and can output to community standard data structures (e.g. AnnData and SpatialData formats)^24,25^.

More specifically, the Sainsc workflow comprises as follows; first, data is read into a custom Rust data structure (a hash map of sparse matrices of equal shape). The reading function supports popular file formats and creates a pixel structure representing the native resolution of the *in-situ* capture SRT data. A sparse data format was chosen to allow for memory efficient handling of sparse *in-situ* capture SRT data. To support imaging-based SRT or unsupported file formats, the Rust data structure can be constructed by providing a Python DataFrame. The optional next step involves filtering genes by count and visualising gene expression over the tissue to select the region of interest for analysis. Then we model the gene expression probability density over the tissue by applying the Kernel Density Estimation (KDE) over all selected genes in Rust. Here, a kernel function (Sainsc has built-in support for a Gaussian and Epanechnikov) and the kernel bandwidth (in pixels or micrometres) is defined and the kernel pre-computed once, stored and subsequently applied as required. To speed up computation and prevent gene expression signal leakage, the Gaussian kernel is cropped at twice the bandwidth (by default). After the KDE is applied, the user can view the probability density of total gene expression over the tissue. In the final step, the cell-type map is generated. This step requires a cell-type-specific gene expression signature matrix, which can either be provided by the user (e.g. from existing single-cell RNAseq data) or computed *de novo* from the data. *De novo* computation of cell-type signatures from the SRT data follows a similar procedure to SSAM, where local maxima of gene expression are considered representative of cell gene expression profiles and clustered to identify cell-type-specific signatures^19^. To minimise the memory and runtime requirements for the pixel-level cell-type assignment, only the subset of genes defined in the signature matrix are used, and small windows (by default, 500 x 500 with a padding according to the kernel radius) are analysed in parallel. Sainsc uses the cosine similarity to identify the most similar cell-type signature for cell-type assignment to each element^26^, which is then rendered as a coloured pixel on the cell-type map.

Cell-type classification often entails variable class resolutions. For instance, certain classes like neural endothelial cells are well-defined with distinct transcriptional profiles. In contrast, other classes, such as various inhibitory or cortical-pyramidal subtypes, require a much finer class resolution within the dataset. Therefore, the similarity between signatures may vary strongly. To effectively account for this, we have adapted a normalization strategy for our standard cosine similarity score. This adapted ‘assignment score’ better reflects the local class resolution by scaling the difference in cosine similarity between the two best scoring cell types according to an upper bound based on the similarity between the two best signatures themselves for every pixel.

**Figure 1.**
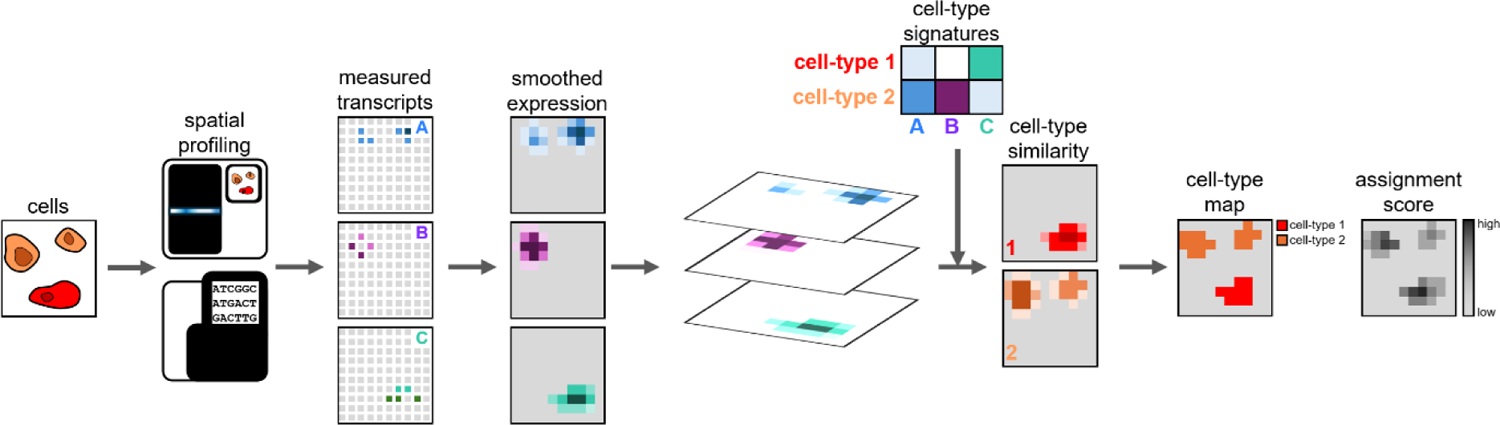
Sainsc workflow. Sainsc takes gene counts from spatially resolved transcriptomics platforms as input and models 2D gene expression using Kernel Density Estimation (KDE). Using reference cell-type signatures (either from prior knowledge or calculated *de novo* from the data), it models cell-type assignment using cosine similarity of the gene expression with the reference signatures. Each spot gets assigned a cell type and an assignment score is calculated to estimate the confidence of the assignment.

To investigate the accuracy of spatial cell-type assignment and assignment score we applied Sainsc to a synthetic dataset^27^. (**Figure 2**). The simulated data consists of 100 replicates of 500 x 500 grids, containing 625 cells each assigned one of nine cell types of the mouse kidney. Spots were assigned gene expression based on the cell identity of the spot using a Poisson process with parameters inferred from single-cell RNAseq data, and zero inflation introduced through uniformly at random setting 20% of spots to zero counts. Sainsc demonstrated comparable accuracy to TopACT (both run at native resolution) and outperformed robust cell type decomposition (RCTD) run with a bin size of 20 (**Figure 2A, B**)^28^. Investigating the effect of filtering spots based on assignment score revealed that the spots with the above median assignment scores had 0.995 accuracy for all but one cell type (**Figure 2C**). Inspecting spatial maps of the assignment score revealed distributions that visually corresponded to a realistic representation of cellular architecture given the ground truth setting (**Figure 2D-G**).

**Figure 2.**
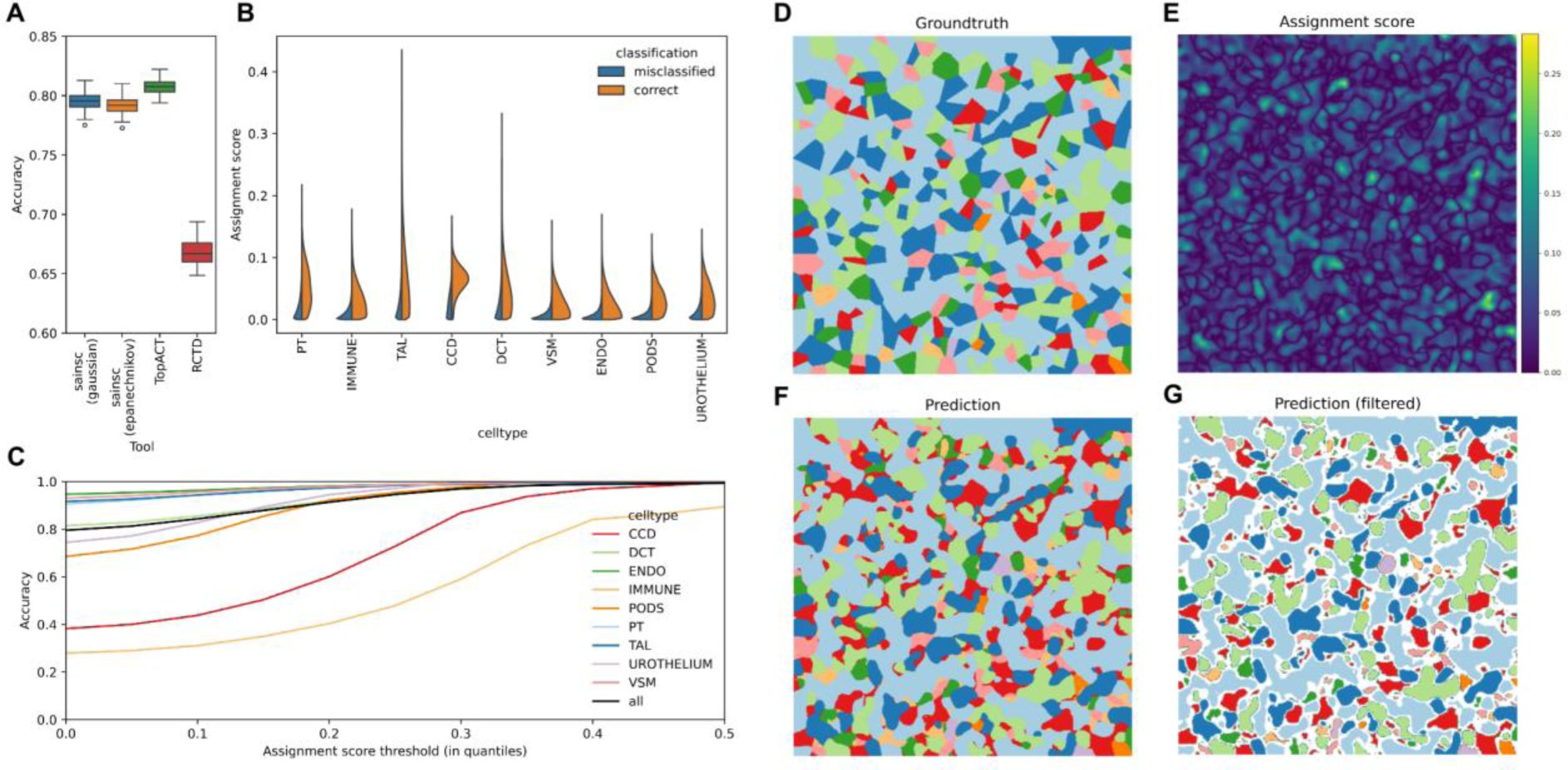
Sainsc cell-type assignments are accurate and increasing assignment score associates with increasing accuracy. (A) Box plot of accuracy of cell-type assignments in 100 simulated samples. From left to right: Sainsc using a Gaussian kernel (bandwidth of 5, truncated at 2 bandwidths), Sainsc with an Epanechnikov kernel (bandwidth of 10), TopACT, and RCTD. Center line, median; box limits, upper and lower quartiles; whiskers, 1.5x interquartile range; points, outliers. TopACT and RCTD results are taken from Benjamin *et al* 2024^27^. (B) Violin plots of cell-type-specific assignment confidence scores in correct and misclassified spots for Sainsc with Gaussian kernel. Misclassified spots show very low assignment scores compared to correctly assigned spots. (C) Line plot of change in Sainsc accuracy when filtering the bottom percentiles of assignment scores. Visualisation of one of the replicates of cell-type map for (D) the ground truth, (E) assignment score map, (F) Sainsc, (G) and Sainsc filtered for bottom 20^th^ percentile of assignments per cell type. CCD, cortical collecting duct; DCT, distal convoluted tubule; ECS, endothelial cells; PT, proximal tubule; TAL, thick ascending limb of the loop of Henle; VSM, vascular smooth muscle.

### Sainsc reconstructs accurate cell-type maps of mouse whole embryo and mouse brain at sub-micron resolution

To investigate the usability of Sainsc with challenging real-world data we applied it to a section of a E16.5 whole mouse embryo data profiled using Stereo-seq^14^. The sample has 28,633 genes detected over 10,582,619,673 reads generated over a large field of view (approximately 1.4 x 0.9 cm), representing one of the largest *in-situ* capture SRT datasets. We used Sainsc to perform both unsupervised and supervised analysis in the native resolution and compared the output to previously published cell-type maps (**Figure 3**). Supervised analysis resulted in highly similar cell-type maps (**Figure 3A, B**). We observed increased sensitivity across the section, including the choroid plexus cells at the fourth ventricle, which was further validated by its marker gene *Ttr* (**Figure 3D, E**). We believe that this increased sensitivity is a direct consequence of the segmentation-free analysis of gene expression in Sainsc. Most notably, Sainsc was able to identify many more erythrocytes compared to the original study, with highest accumulation of signal in expected organs: the spleen, liver and umbilical cord. Interestingly, we found erythrocytes in the choroid plexus at the fourth ventricle (**Figure 3D**). We have previously shown that current cell-segmentation algorithms struggle with segmenting choroid plexus cells^19^. The choroid plexus produces and secretes the majority of the cerebral spinal fluid and consists of ependymal cells surrounding a core of capillaries^29^, rationalising Sainsc’s detection of erythrocytes adjacent to ependymal cells. Prediction of erythrocytes by Sainsc was supported by expression of erythrocyte marker gene encoding for haemoglobin beta adult S chain (*Hbb-bs*) (**Figure 3F**).

While using existing cell-type-specific signatures can be a convenient way to transfer cell-type labels, inherent differences between profiling technologies may warrant the need for unsupervised analysis. Here, Sainsc adopts an unsupervised clustering approach to generate cell-type signatures from the tissue itself in the same way as the SSAM algorithm^19^. Briefly, clustering is performed on a subset of pixels which correspond to local maxima of gene expression, under the assumption that regions of high expression are likely to originate from within cells. The unsupervised analysis of the mouse embryo section identified 33 clusters that corresponded to the expected organ structure (**Figure 3C**). The unsupervised analysis also resulted in superior detection of erythrocytes in the spleen and liver (**Figure 3G-L**). Erythrocytes are hard to identify in a DAPI segmentation-based analysis due to the lack of a nucleus in mature erythrocytes. We were also able to correctly distinguish ossifying regions (as demonstrated by expression of *Col1a1* and *Col1a2*)^30^ from other chondrocytes (expression of marker genes *Sox9* and *Col2a1*)^31^ (**Figure 3M-R**) as shown by unsupervised analysis using FICTURE on the same dataset^32^.

**Figure 3.**
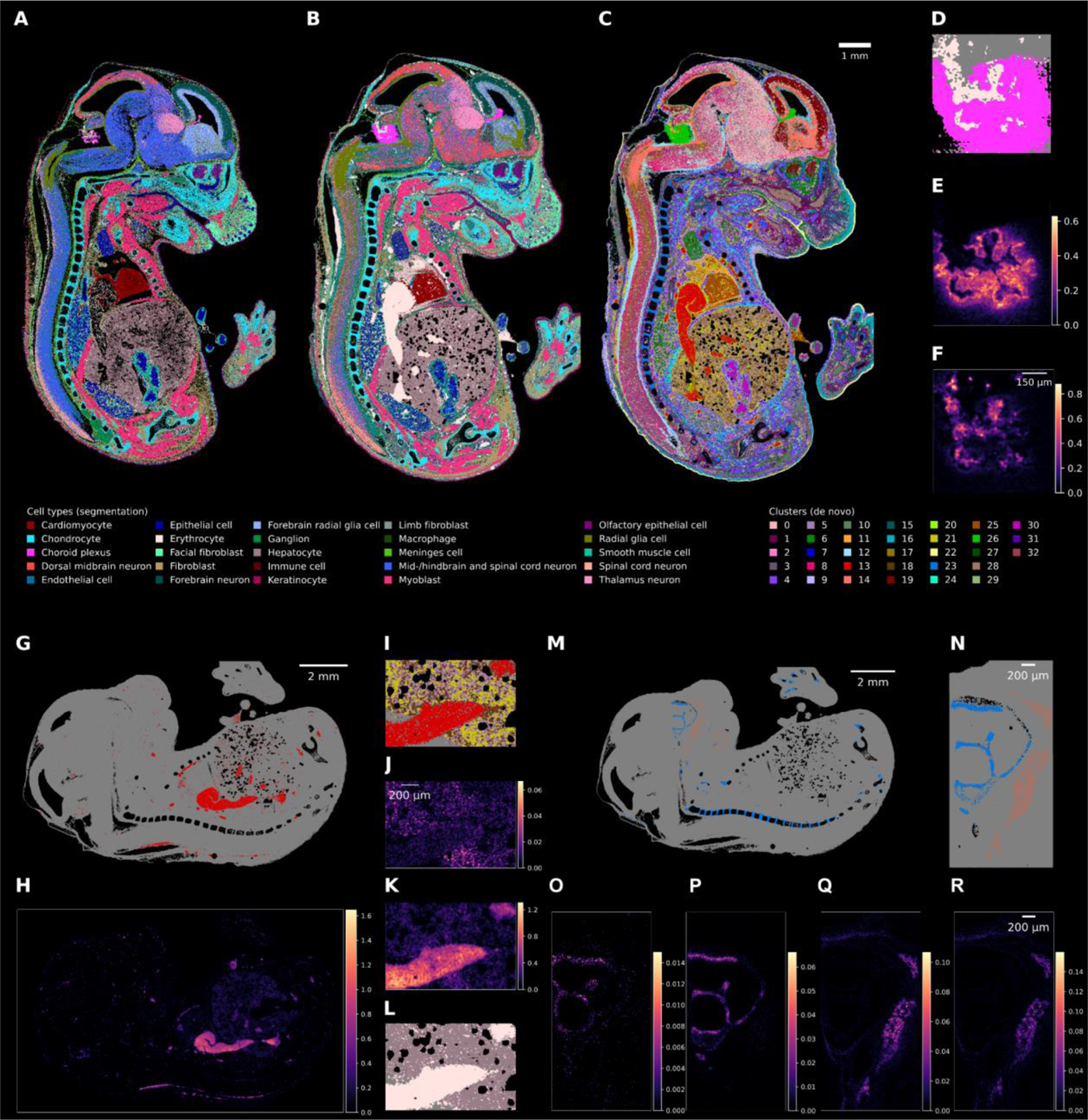
Sainsc enables whole organism (mouse embryo) cell type assignment at subcellular resolution. (A) Segmentation-based cell-type assignment from Chen *et al*^14^ (B) supervised cell-type map using Sainsc with cell-type signatures extracted from the segmentation-based data and (C) unsupervised cell-type map using Sainsc with *de novo* signatures. (D-F) Choroid plexus region of the fourth ventricle for Sainsc supervised cell type assignment showing (D) the structuring of the choroid plexus and its supply with erythrocytes highlighted by the expression of marker genes (E) *Ttr* (choroid plexus) and (F) *Hbb-bs* (erythrocytes). (G-L) Distribution of erythrocytes throughout the embryo (G) in the unsupervised approach (cluster 13) validated by (H) the erythrocyte marker gene *Hbb-bs*. (I-L) Region showing parts of the spleen and liver for (I) the unsupervised approach, the expression of (J) hepatocyte marker albumin (*Alb*), and (K) erythrocyte marker *Hbb-bs*, and (L) erythrocytes and hepatocytes in the supervised cell-type assignment (M, N) Assignment of Cluster 23 and 28 correspond to chondrocytes and ossification activity, respectively as validated by the expression (KDE) of (O) *Sox9*, (P) *Col2a1*, (Q) *Col1a1*, and (R) *Col1a2*.

To further demonstrate the performance of Sainsc in delineating complex cell-type organisation, we applied it to mouse brain samples. First, we analysed a Stereo-seq dataset of a mouse brain coronal section. Performing supervised analysis using cell-type signatures derived from the segmentation-based analysis of the same tissue. We were able to identify all major anatomical regions of the brain (**Figure 4A, B**). Furthermore, we identify similar distributions of cell-types in another mouse brain coronal section profiled by another full-transcriptome sub-micron resolution *in-situ* capture SRT technology, Nova-ST (**Figure 4C**)^17^. To further investigate the intricate cellular patterns, we generated a cell-type map using signatures from Yao *et al* 2021^33^, which has cell-type-specific gene expression signatures for layer-specific neurons in the isocortex and hippocampal formation regions of the cortical plate. The cell-type map finely resolved the locations of numerous layer and region-specific neurons as well as the fine lining of vascular and leptomeningeal cells (VLMC) at the border of the tissue (**Figure 4D-Q**). This demonstrates that Sainsc can leverage cell-type signatures obtained by different technologies, such as single-cell RNAseq (scRNAseq) or other SRT technologies, to generate the subcellular cell-type classification.

Prior studies have highlighted the difficulty of finding rare cells such as macrophages in the central nervous system^27^. To investigate the localisation of border-associated macrophages (BAMs), we annotated spots using the cell-type signatures from Yao *et al* 2023^34^. The identified BAMs were distributed in the in the expected meningeal and perivascular spaces regions (**Figure 4R**)^35^. In particular, we identified BAM localising to the VLMC border layer, that were not reported in a previous characterisation of macrophage distribution in the same dataset^27^. This localisation of BAMs is consistent with coronal sections of the mouse brain profiled using MERFISH^36^.

**Figure 4.**
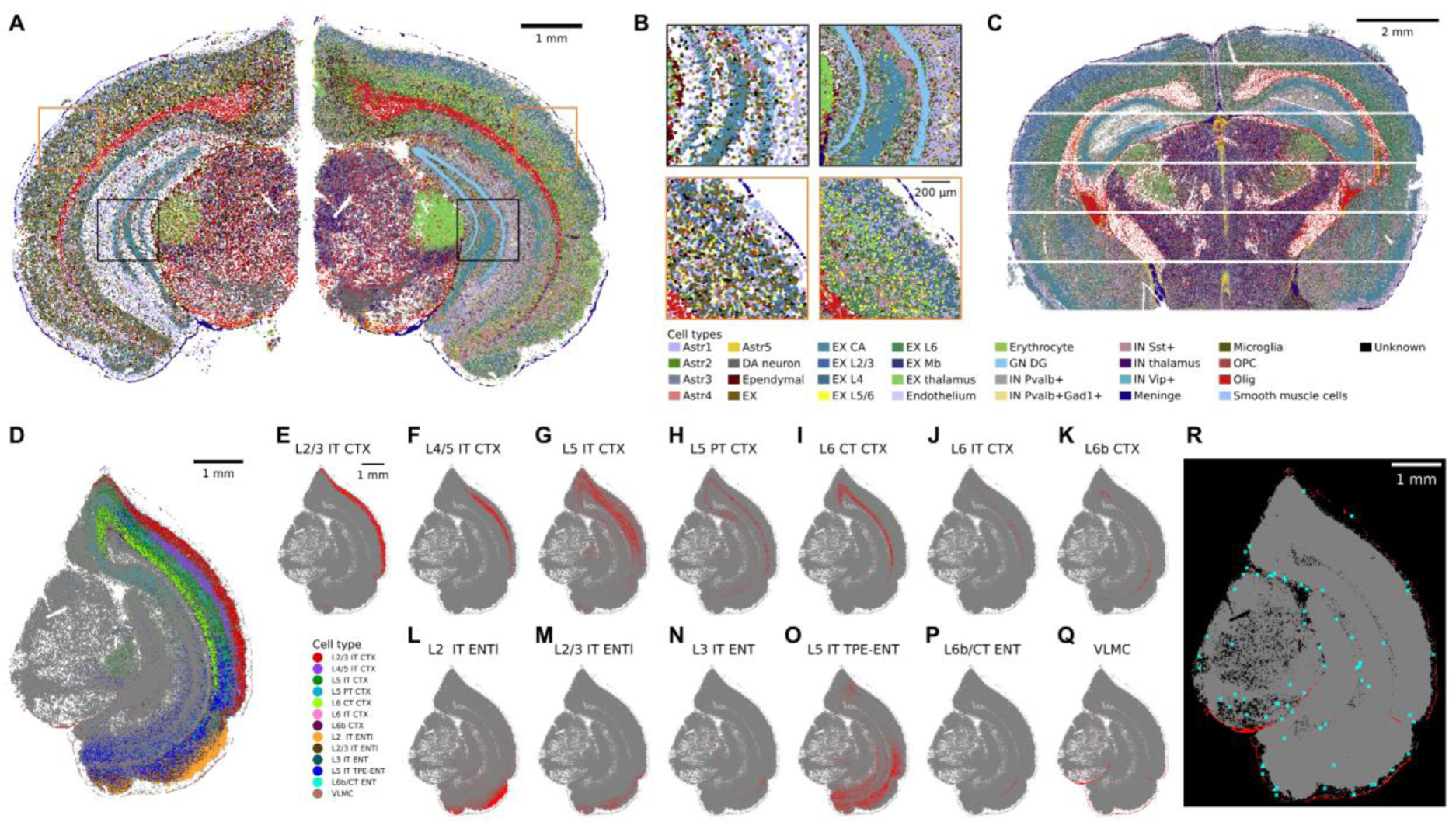
Sainsc accurately resolves cortical layering in Stereo-seq data and identifies rare cell types in mouse brain sections. (A, B) Comparison of segmentation-based (left) and Sainsc (right) cell-type maps of a coronal section of the mouse brain profiled by Stereo-seq. (C) Cell-type signatures are transferrable to different technologies as demonstrated by cell-type assignment in a coronal brain section profiled with Nova-ST. (D-Q) Using scRNAseq based signatures Sainsc resolves the intricate structures of the cortical plate as shown for (E-P) the neuronal layers of the cortical plate and (Q) the surrounding VLMC. (R) Localisation of identified border-associated macrophages (cyan) is close to meningeal and vascular cells (red). Vascular subclasses include ABC NN, VLMC NN, Peri NN, Endo NN, and SMC NN. ABC, arachnoid barrier cells; CTX, isocortex; Endo, endothelial cells; ENT, entorhinal area, ENTl, lateral entorhinal area; IT, intratelencephalic; NN, non-neuronal; Peri, pericytes; PT, pyramidal tract; SMC, smooth muscle cells; TPE, temporal association, perirhinal, and ectorhinal areas; VLMC, vascular leptomeningeal cells.

### Sainsc is applicable to imaging-based SRT

Imaging-based SRT platforms with high sensitivity and larger gene panel are increasingly becoming commercially available. To demonstrate the applicability of Sainsc to these methods, we used it to analyse a publicly available Xenium dataset of a coronal section of the mouse brain (**Figure 5**). We compared the cell-type map produced by Sainsc to cell types annotated in a previous study that used differentially expressed marker genes from scRNAseq data (**Figure 5A, B**)^20,34,36^. To ensure comparability, the gene expressions signatures were calculated from the segmented and annotated Xenium data and used for supervised cell-type annotation with Sainsc. The resultant cell-type maps showed remarkable similarity; however, we observed several differences. We observed inconsistent annotation of the inhibitory neurons classified as “RT ZI Gaba”, where Sainsc annotated two distinct domains with this cluster label, one of which was not annotated in the prior study (**Figure 5D-G**). The class annotates GABAergic inhibitory neurons from the thalamic reticular nucleus (RT) and the zona incerta (ZI) neurons in hypothalamus, that have a shared developmental origin^34,37^. Referring to the Allen Brain Atlas, we indeed found that the RT and ZI domains were spatially distinct (originating from the thalamus and hypothalamus, respectively)^38^, and matched the annotated regions by Sainsc (**Figure 5E, L**). In the previous annotation, the cells in the RT were annotated correctly, however neurons in the ZI were annotated less precisely as “inhibitory GABA mixed 2” (**Figure 5D**).

Furthermore, we observed differences in the thalamus. In particular, a region corresponding to the paraventricular nucleus of the thalamus (PVT), was annotated to “MEA-COA-BMA Slc17a6 Glut” by Sainsc, compared to the rest of the region which was annotated as thalamic glutamatergic neurons (“Thal Glut Neurons”, **Figure 5H, I**). The gene expression signatures of “MEA-COA-BMA Slc17a6 Glut” and “Thal Glut Neurons” are highly similar (cosine similarity of 0.79). The assignment score in the PVT region was much lower than the rest of the thalamus (**Figure 5C, J**), which lead us to suspect that the combination of the cell-type signature matrix and Xenium gene panel was not optimal for annotating pixels in the PVT region. While it is expected that the PVT should be enriched for thalamic glutamatergic neurons, it has been shown that the PVT has specialised glutamatergic neuronal subclass (“PVT-PT Ntrk1 Glut”) highlighted by the expression of one of its marker genes *Bcl11b* (**Figure 5K**)^34^. However, this subclass was not present in the gene expression signature matrix used to annotate the sample. We hypothesised that the lack of a PVT-PT specific signature led to the misclassification by Sainsc with low assignment scores. To investigate whether this hypothesis was true, we revised cell-type assignment using a signature matrix with the class annotations of Yao *et al* 2023 (34 classes)^34^, also including the “PVT-PT Ntrk1 Glut” subclass. Indeed, we observed that pixels in the PVT region were now annotated correctly and with a high assignment score (**Figure 5M, N**).

**Figure 5.**
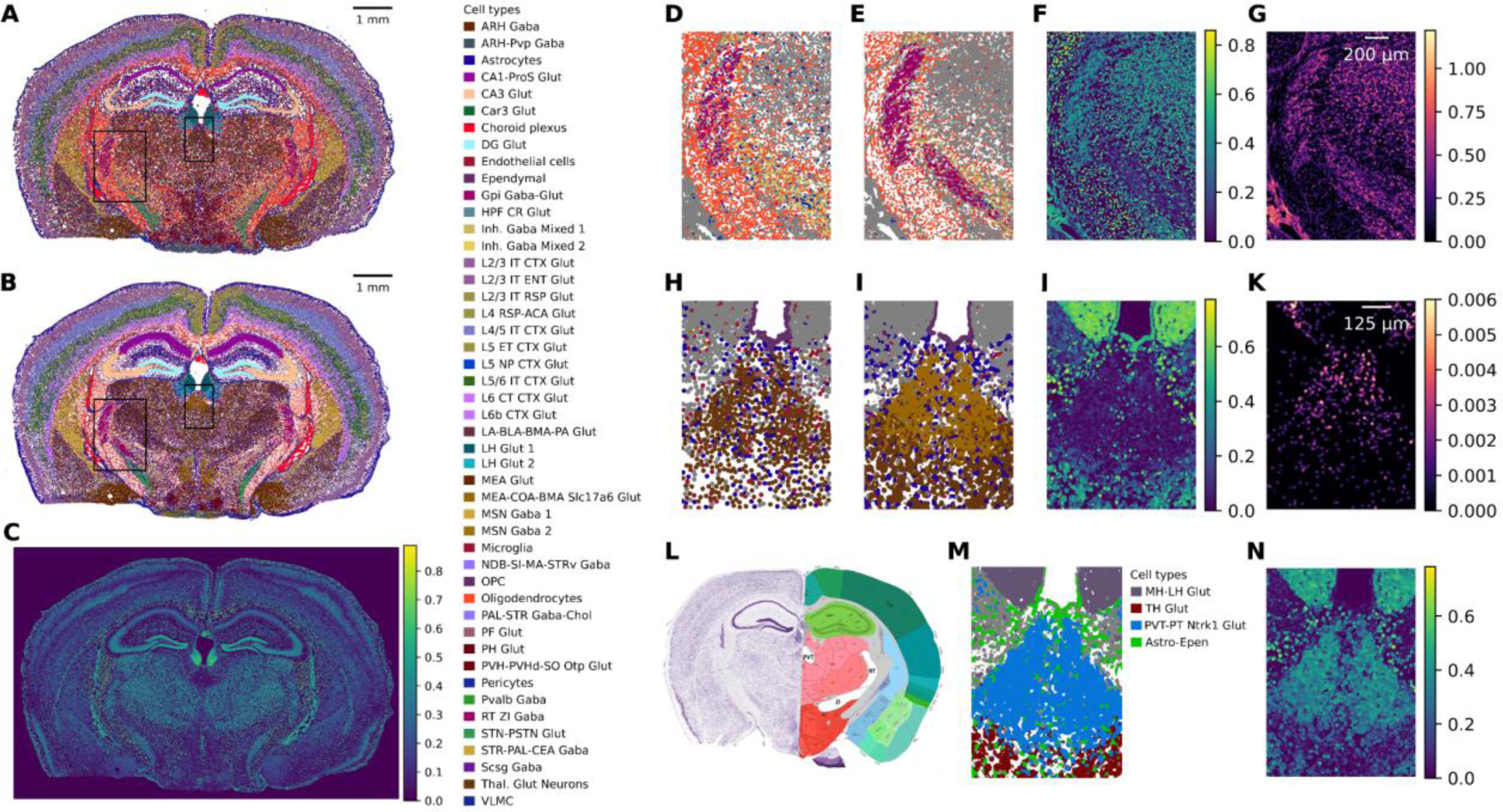
Sainsc is applicable to Xenium and the assignment score guides identification of unrepresented cell types and regions. (A) Reference cell-type map based on segmentation, (B) Sainsc’s supervised cell-type assignment and (C) assignment score of a Xenium coronal mouse brain section. (D-G) Zoom in of the RT and ZI regions showing (D) the original cell-type assignments, (E) Sainsc cell-type assignments, (F) Sainsc assignment score, and (G) KDE of total gene expression. (H-K) Misassigned cell type in the paraventricular nucleus of the thalamus (PVT), showing the (H) the original cell-type assignments, (I) Sainsc cell-type assignments, (J) Sainsc assignment score, and (J) KDE of marker gene for PVT neurons, *Bcl11b*. (L) Anatomical annotations of a similar section of the mouse brain from the Allen Mouse Brain Atlas and Allen Reference Atlas, mouse.brain-map.org (https://atlas.brain-map.org/atlas?atlas=1&plate=100960236, P56, Coronal section 72 of 132)^39–42^.

### Computational performance

A major problem associated with the newest generation of *in-situ* capture SRT methods is handling and processing the large volumes of data generated – the combination of high spatial resolution and full transcriptome coverage increases processing time and memory requirements. Efficient data processing, however, is important to enable exploratory analysis, e.g. through interactively modifying parameters to improve the output. We measured Sainsc runtime and memory usage for varying number of genes and cell-type signatures when analysing the Stereo-seq mouse hemibrain section with 8 threads. The increase in wall time and CPU time scales linearly with both the number of genes and cell-types used in the analysis (**Figure 6A, B**). While the memory usage also depends on these two parameters, its increase is moderate (~4 GB for 250 genes and 10 cell types and < 6.5 GB for 16,000 genes and 40 cell types) (**Figure 6C**). Often, only a few thousand highly/spatially variable genes are used for clustering and cell-type assignment. Sainsc’s wall time for a typical run (e.g. 2,000 genes and 20 cell types) is below 10 min, including data loading, therefore effectively enabling interactive exploration of the data.

**Figure 6.**
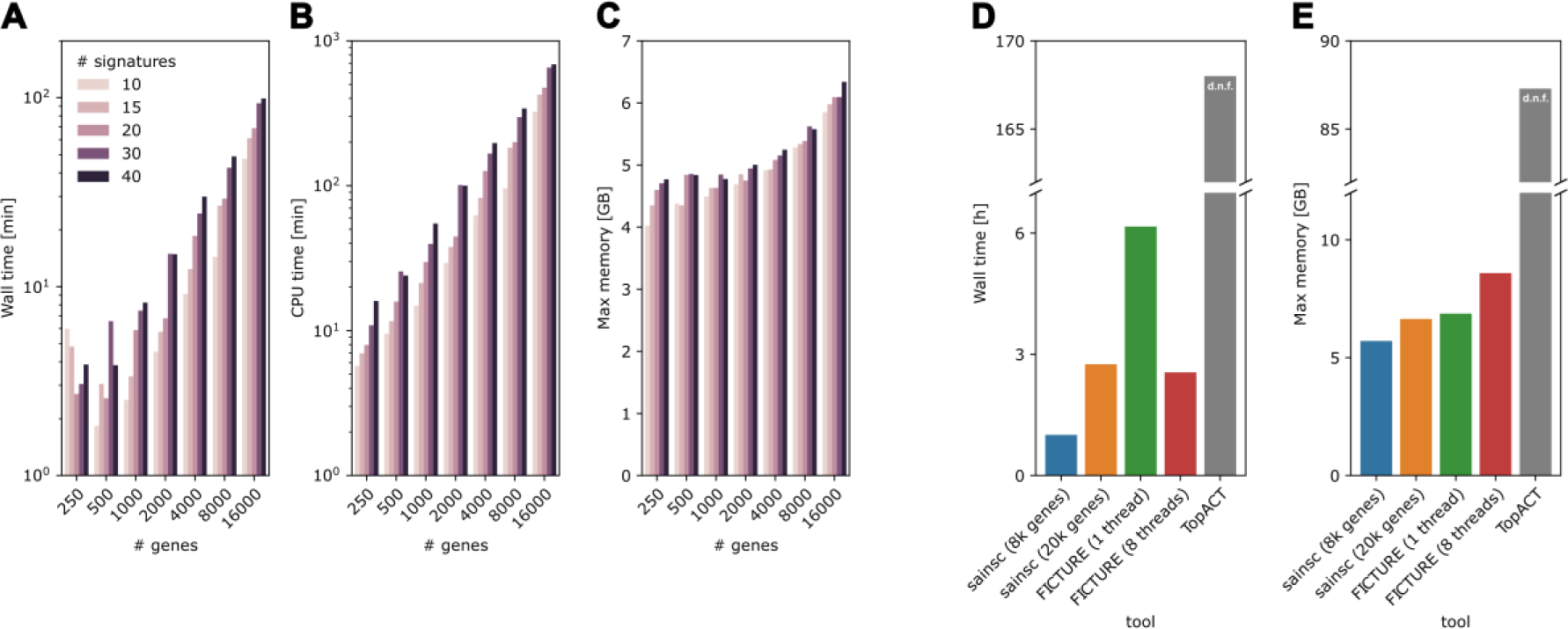
Sainsc is scalable and has low computational resource requirements. (A) Wall time, (B) CPU time, and (C) maximum memory usage when analysing Stereo-seq coronal hemibrain data with varying number of genes and cell types. Comparison of (D) runtime and (E) maximum memory usage of Sainsc, TopACT, and FICTURE. d.n.f.; did not finish.

Two recent computational tools, FICTURE and TopACT^27,32^, also demonstrate accurate performance for creating cell-type maps for SRT data. FICTURE claims to be efficient, while TopACT does not report processing time or memory requirements. We compared the computational performance of Sainsc against FICTURE and TopACT applied to the Stereo-seq mouse brain sample (**Figure 6D, E**). To ensure comparability between FICTURE and Sainsc we ran FICTURE using one core and 8 cores (as used by Sainsc), and Sainsc with 7,972 genes and 19,708 genes (as used by FICTURE). We recorded wall time and maximum memory used in processing the Stereo-seq adult mouse coronal hemibrain sample (**Figure 6D, E**). For all metrics, Sainsc performed similar to FICTURE and substantially better than TopACT. In multi-threaded mode Sainsc and FICTURE wall times were approximately equivalent at less than 3 hours. TopACT running with 8 processes did not finish analysis in the allotted time and the job was cancelled after 7 days. Furthermore, Sainsc required had similar maximum memory usage to FICTURE in single-threaded mode. The reduced runtime of FICTURE in multi-threaded mode compared to its single-threaded equivalent comes at a cost of increased memory usage however, approximately 30% more than Sainsc. TopACT used substantially more memory (~87 GB of RAM) during its runtime than either Sainsc or FICTURE.

Julia has been described to have the interactivity and syntax of scripting languages with the speed of compiled languages^43^. As such, we also implemented the core functionality of Sainsc into a Julia package, Sainsc.jl. To demonstrate the correspondence of the Julia with the Python/Rust implementation we analysed the Stereo-seq mouse coronal hemibrain section with signatures obtained from segmented data and achieved near identical results (discordantly classified pixels: 9.2 x 10^-5^%; median absolute difference of assigned pixels for the assignment score: 7.5 x 10^-9^, for the cosine similarity: 0, and for the total mRNA KDE: 2.4 x 10^-7^) while utilising similar computational resources (**Figure 7**).

**Figure 7:**
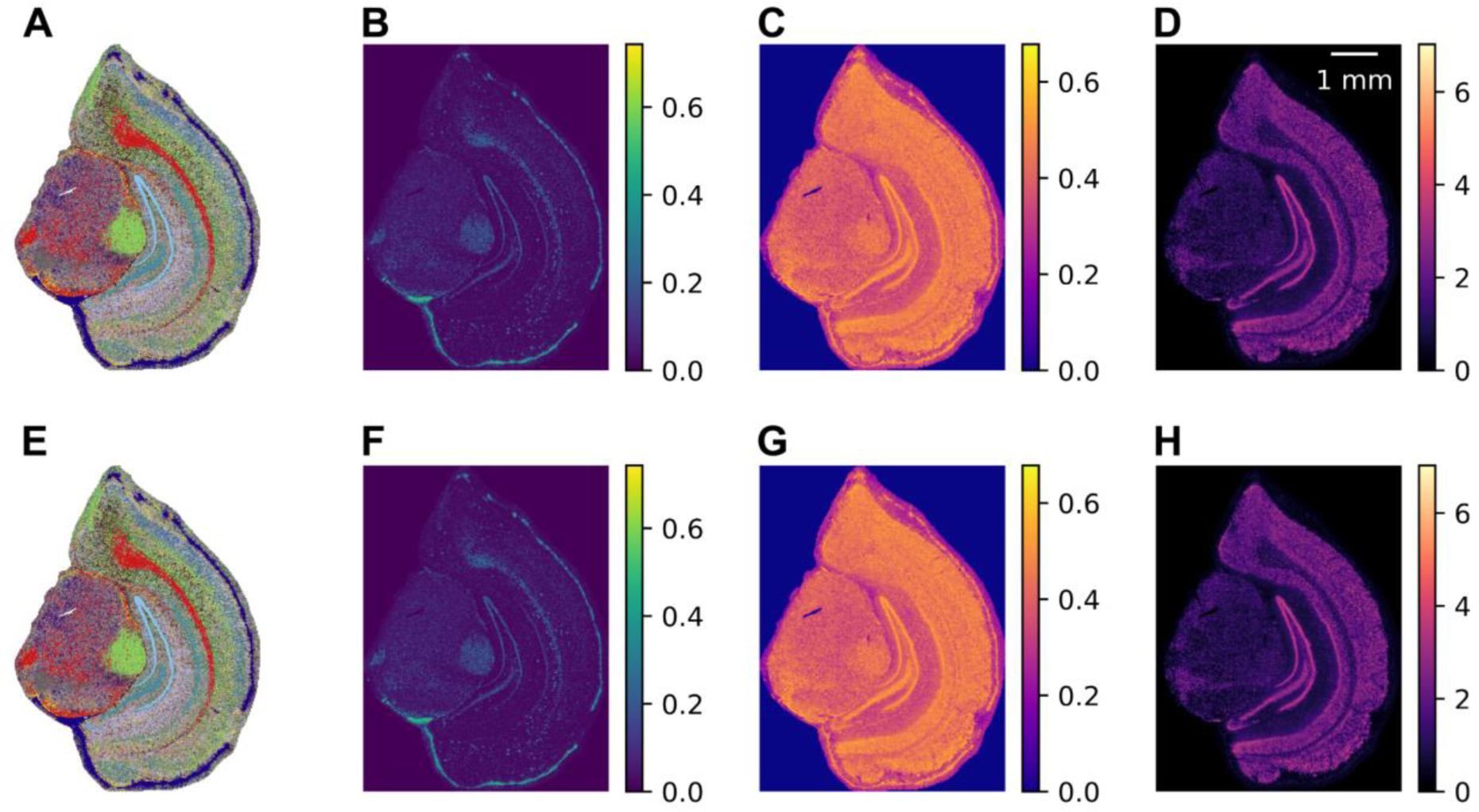
Sainsc’s Python and Julia implementation generate near identical results. Comparison of the Stereo-seq mouse hemibrain when analysed with (A-D) Sainsc (Python) and (E-H) Sainsc.jl (Julia). Shown are (A, E) the unfiltered cell-type map, (B, F) the assignment score, (C, G) the cosine similarity of the assigned cell type, and (D, H) the KDE of the total mRNA.

## Discussion

The most recent wave of high-resolution full-transcriptome *in-situ* capture SRT methods are an attractive choice for characterising cellular heterogeneity in tissue sections, however, to date there is a lack of performant methods to analyse these types of data. Furthermore, as there is no “one size fits all” workflow, analysis needs to be exploratory and interactive, to e.g. optimise the clustering parameters to obtain the correct number of clusters or testing the efficacy of annotating spatial gene expression from signature sets obtained from different single-cell studies. Such an interactive analysis paradigm demands fast and performant algorithms with minimum computational requirements.

In this study we present a novel computational tool, Sainsc. Applying Sainsc to simulated data, we achieve top tier accuracy for classification and show that our cell-type assignment score can be used as a filter to improve the confidence of the results. We then accurately map the cellular landscape at the whole organism level, improving on previous segmentation-based analysis. We demonstrate that Sainsc was the most computationally performant when analysing the mouse brain requiring only ~1 hour of CPU wall time and ~5 Gb of memory. The moderate resource requirements of our implementation will facilitate exploratory analysis of extremely large datasets. Furthermore, Sainsc’s annotations can easily scale up to several hundred cell types by integrating the expression landscape of thousands of genes.

We demonstrate the additional utility of the assignment score for interpreting the quality of the results. While annotating cell-types based on gene expression signatures obtained from single-cell RNAseq is a quick way to contextualise new data in the light of prior knowledge, we show that when the gene panels or cell type definitions differ from the reference source, it can result in lower confidence cell-type assignment. To the best of our knowledge, we are the first to demonstrate the value of a visual map of the assignment score to effectively evaluate and improving assignment confidence at the organ and organism scale.

We also compared Sainsc to two other existing tools in this space, TopACT and FICTURE. While TopACT was highly accurate, its computational inefficiency makes it unsuitable for exploratory analysis. We were unable to complete analysis of the mouse coronal hemibrain Stereo-seq sample with TopACT in under seven days. FICTURE performed similar to Sainsc on the same gene set, however, in our analysis we noticed that if chosen appropriately, much fewer genes are necessary to distinguish cell types accurately (e.g. for the Xenium dataset 248 genes are sufficient to separate 47 cell types). This reduced number of genes comes at great performance benefits for Sainsc especially in terms of runtime. Taken together, we believe that Sainsc is currently the only computational tool to analyse such large datasets accurately with resource requirements that approach the scope of consumer electronics. Further emphasis is placed on efficiency when considering the spatial genomics field is going: towards 3D cell atlas^44^, multi-omics^45^ and entire human organs^46^. We believe Sainsc is uniquely suited to be further developed to cater for the even higher computational needs of increasing complex experiments without sacrificing accuracy.

## Methods

All analyses were performed in Python (v3.10.14) with Sainsc (v0.0.1, compiled using rustc v1.76.0) and the following main packages; pandas (v2.2.2), polars (v0.20.31), scipy (v1.14.0), numpy (v1.26.4), scanpy (v1.10.2), squidpy (v1.5.0), anndata (v0.10.8).

### Sainsc

#### Gene expression estimation

Sainsc adopts the estimation of gene expression in 2D based on kernel density estimation from SSAM^19^. In short, the gene expression density *p̂* is estimated as

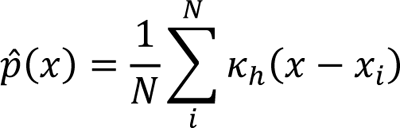

where *k*_*h*_ is the kernel function in dependence of the bandwidth *h*, *N* the number of data points, and *x_i_* the location of the *i*th spot of the given gene. As simplification, however, we only consider a discrete space (pixels) for both the gene expression and the estimated probability distribution which allows for pre-computed kernel “masks” and efficient calculation.

#### Cell-type assignment

Cell-type assignment to each spot (pixel) is based on cosine similarity. The cosine similarity measures the similarity of two vectors via their angle and can be efficiently computed using the inner product.

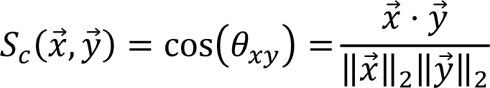

The cosine similarity is calculated between a pixel and each cell-type signature, and the most similar cell type is assigned.

### Assignment score

Let 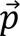 be the gene expression of a pixel (for all genes under consideration) and 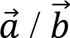 the signatures of the most and 2^nd^ most similar cell types, respectively. We define the assignment score as

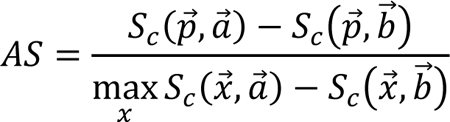

where max 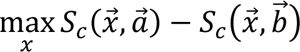 is the upper bound of the difference in cosine similarities given 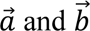. Assuming non-negative gene expression the following must hold 0° ≤ *θ_pa_* ≤ *θ_pa_* ≤ 90°. For simplicity we assume normalized cell-type signatures i.e. 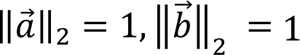.

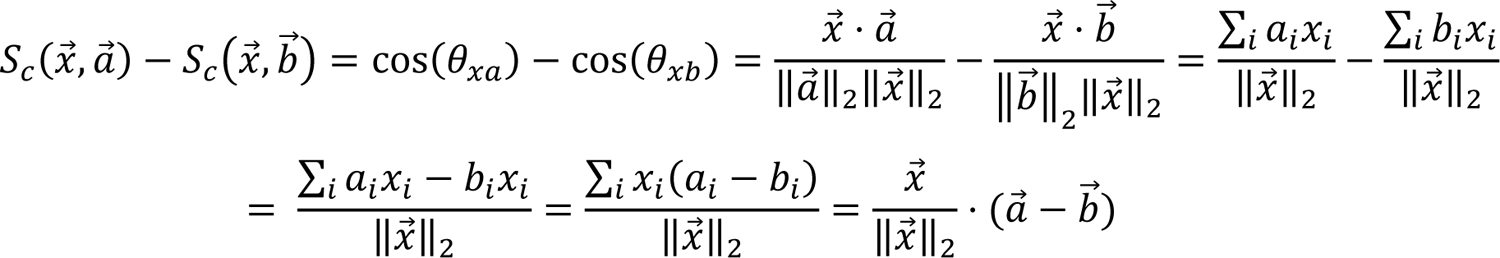

The inner product of 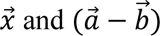 is maximal for colinear vectors and therefore 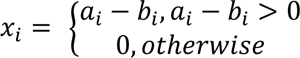 (where negative dimensions are set to 0 due to the non-negativity constraint).

### Sainsc.jl

Conceptually, the Julia implementation is equivalent to the Python/Rust implementation and most data structures (e.g. hashmap of same sized sparse matrices for counts) and processing steps (e.g. processing the data in chunks when generating the cell-type map) are “identical”. As the ecosystem is still young and evolving, Sainsc.jl integrates with multiple single cell/spatial transcriptomics frameworks, including Muon.jl which allows writing the gene expression of local maxima to h5ad files which can then be easily read from Python.

To compare the Julia with the Python/Rust implementation of Sainsc we analysed the Stereo-seq hemibrain dataset in Julia (v1.10.4) with Sainsc.jl (commit ea4cfdf031d937da783cf55351a33d693d4a5618) using the same parameters as described with the segmentation-based cell-type signatures. Results were plotted in Python to make them visually comparable.

### Synthetic data

To compare Sainsc to TopACT we used the simulated data from the TopACT publication^27^. For Sainsc the bandwidth was set to 5 for the Gaussian kernel (truncated at 2x bandwidth) and to 10 for the Epanechnikov kernel. The average cell-type expression was used as cell-type signatures for Sainsc and the accuracy was calculated using *sklearn.metrics.accuracy_score*. Results for TopACT and RCTD were taken from the publication.

### Stereo-seq analysis

All Stereo-seq datasets were analysed at native resolution with 500 nm / px (equivalent to the 500 nm bead-to-bead distance). A Gaussian kernel with bandwidth 8 px (4 μm) truncated at 2 bandwidths was used throughout.

#### Embryo

The E16.5 E1S3 embryo data from the original Stereo-seq publication was cropped and masked to the region of interest. The cell-type assignment was performed and for visualization the background was filtered based on cell-type-specific and global thresholds for the KDE of the total mRNA. For the supervised analysis signatures were extracted from the segmented data provided in the original publication by selecting the top 2,000 highly variable genes (HVG) (*scanpy.pp.highly_variable* with flavor “seurat_v3”) and averaging across all cells per cell type.

#### Embryo Unsupervised

For the *de novo* analysis the data was read as 50x50 bins and filtered for bins (“cells”) with more than 1,000 counts and genes with more than 100 counts. The top 10,000 HVG were identified using *scanpy.pp.highly_variable* with flavor “seurat_v3”. A spatial neighbor graph was build using *squidpy.gr.spatial_neighbors* (coord_type=”grid”, n_rings=1, n_neighs=4) and the top 2,000 spatially-variable genes by Moran’s I of the 10,000 HVGs were identified using *squidpy.gr.spatial_autocorr* (with mode =”moran”). Next, the full resolution data was loaded with Sainsc and the local maxima identified (s*ainsc.LazyKDE.find_local_maxima* with min_dist=4 and min_area=60). The local maxima were loaded as AnnData object using the previously identified SVGs, the counts per “cell” normalized (*scanpy.pp.normalize_total*) and log-transformed (scanpy.pp.log1p). The data was PCA transformed (*scanpy.pp.pca*) and the neighborhood graph, based on the latent dimensions, constructed (*scanpy.pp.neighbors*). The data was then clustered using Leiden community detection (*scanpy.tl.leiden* with resolution=2). The “cell-type” signatures were created as average per cluster and used for cell-type assignment.

#### Coronal mouse hemibrain

To compare Sainsc to the segmentation-based cell type assignment of the mouse coronal hemibrain section, signatures were extracted as average per cell type for the differentially expressed genes (7,972) reported in Yao *et al* 2023^34^. To demonstrate the efficiency in identifying the isocortex layers cell-type signatures from scRNA-seq data from Yao *et al* 2021^33^ were calculated per “subclass” (downsampled to 5,000 cells per subclass) for the differentially expressed genes reported in the same study.

#### Border-associated macrophage detection

To detect border-associated macrophages (BAM), we generated cell-type signatures from the Allen brain atlas (10x Genomics Chromium v2 data) from Yao *et al* 2023^34^ for all subclasses (down sampled to max. 500 cells per subclass) resulting in 279 signatures. Using the differentially expressed genes from the same publication the Stereo-seq brain was annotated as previously described. The binary map of BAM assigned spots was labelled as connected regions using *skimage.measure.label* and the centroids extracted using *skimage.measure.regionprops_table* and filtered for regions with an area of at least 60 pixels.

### Xenium analysis

The Xenium brain dataset from 10X genomics (replicate 1) was analysed by binning all transcript counts to 500 nm. The cell-type signatures were based on the segmented Xenium dataset by Salas *et al*^20^. A Gaussian kernel with bandwidth 8 px (4 μm) truncated at 2 bandwidths was used.

To correctly assign cell-types in the PVT region we used cell-type signatures generated from downsampled scRNAseq data^34^ based on the “class”-level. Additionally, the “PVT-PT Ntrk1 Glut” subclass was added to the signatures and the analysis repeated as described above.

### Nova-ST analysis

NovaST data was binned to 500 nm and analysed using the signatures based on the Stereo-seq segmented mouse hemibrain slice for the differentially expressed genes defined in Yao *et al* 2023^34^ with a Gaussian kernel of bandwidth 8 px truncated at 2 bandwidths.

### Benchmarking and runtime performance metrics

The Stereo-seq mouse brain sample was used to benchmark the runtime performance of Sainsc against other methods. To calculate runtime performance metrics jobs were submitted using slurm *sbatch* to a single compute node (-N 1) with 8 CPUs (-n 8) with each node consisting of a Dell PowerEdge C6520 with 2x Intel Xeon Gold 6130 or 6252 @ 2.1 GHz. Jobs were assigned 128 Gb of RAM, unless otherwise specified. The metrics were calculated from the slurm job information (using ‘ElapsedRaw’ as wall time, ‘TotalCPU’ as CPU time, and ‘MaxRSS’ as maximum memory usage) for the entire workflow including data loading, pre-processing, and “spot”-wise cell-type/factor assignment.

#### Sainsc

Sainsc was run with 8 threads, and a Gaussian kernel of bandwidth 8 truncated at 2 bandwidths. To compare the scalability with the number of genes and cell types, cell-type signatures from Yao *et al* 2021^33^ and genes appearing in both the data and the signatures were randomly subsampled and submitted with 68 Gb of RAM. For the comparison with FICTURE and TopACT cell-type assignment was performed for the 42 “subclasses” defined in Yao *et al* 2021^33^ and the number of genes for the signatures was varied; 7,972 (differentially expressed genes from Yao *et al* 2023 that are detected in the data), and the 17,275 genes as used by FICTURE after its default filtering.

#### FICTURE

For FICTURE (v0.0.3.1)^32^ the Makefile was generated using the *run_together* command and non-essential processing steps removed (e.g. plotting and bulk differential gene expression) resulting in an analysis consisting of the main steps *make_spatial_minibatch, make_dge, fit_model, transform*, and *slda_decode*. Furthermore, the *sort* commands were adjusted to a buffer-size of 4Gb (-S 4G). The analysis was performed with 42 factors (corresponding to the 42 cell types used for the other tools) and a width of 15 (corresponding to 8.7 μm side-to-side width of the hexagon following the original publication) while keeping other parameters at the default. The analysis was run with 1 thread / 1 job and 8 threads / 2 jobs.

#### TopACT

TopACT (v1.1.0)^27^ was trained on single cell data from Yao *et al* 2021^33^. The data was subsampled to at most 1,000 cells per cell type leaving ~37,000 cells prior to running the analysis. For classification all pixels within the convex hull around the gene expression were considered and the classification was run in parallel on 8 processes for a radius from 3-9 (min_scale and max_scale parameters, respectively) following the original publication.

## Data availability

All data used in this study is publicly available. Stereo-seq mouse brain and embryo data was downloaded from the original Stereo-seq study (https://db.cngb.org/stomics/mosta/).^14^ Xenium mouse brain data was obtained from 10x Genomics (https://www.10xgenomics.com/datasets/fresh-frozen-mouse-brain-replicates-1-standard).

Nova-ST mouse brain data^17^ was downloaded from Gene Expression Omnibus (https://www.ncbi.nlm.nih.gov/geo/query/acc.cgi?acc=GSM8093729). The scRNAseq data for the isocortex and hippocampal formation from Yao et al^33^ was downloaded from the NeMO Archive (https://assets.nemoarchive.org/dat-jb2f34y) and for the whole mouse brain from Yao *et al* 2023^34^ (https://portal.brain-map.org/atlases-and-data/bkp/abc-atlas).

## Code availability

The Sainsc Python package is available at https://github.com/HiDiHlabs/sainsc and the Julia implementation at https://github.com/niklasmueboe/Sainsc.jl. The analysis code to reproduce the results of this study is deposited at https://github.com/HiDiHlabs/sainsc-study and will be made available upon publication.

## Author contributions

N.I. conceived the study. N.M-B implemented Sainsc, Sainsc.jl and performed data analysis and interpretation. S.T. assisted with interpretation of mouse brain results and provided critical comments and feedback. N.I. and N.M-B wrote the manuscript. All authors commented on and critically revised the manuscript.

## Acknowledgements

We thank Alexander Malt for the review and feedback of the sainsc user guide and documentation. This research has received funding from the Federal Ministry of Education and Research of Germany in the framework of SAGE (project number 031L0265).

